# Kinetic investigation reveals an HIV-1 Nef-dependent increase in AP-2 recruitment and productivity at endocytic sites

**DOI:** 10.1101/2023.04.18.537262

**Authors:** Yuichiro Iwamoto, Anna Ye, Cyna Shirazinejad, James H. Hurley, David G. Drubin

## Abstract

Lentiviruses express non-enzymatic accessory proteins whose function is to subvert cellular machinery in the infected host. The HIV-1 accessory protein Nef hijacks clathrin adaptors to degrade or mislocalize host proteins involved in antiviral defenses. Here, we investigate the interaction between Nef and clathrin-mediated endocytosis (CME), a major pathway for membrane protein internalization in mammalian cells, using quantitative live-cell microscopy in genome-edited Jurkat cells. Nef is recruited to CME sites on the plasma membrane, and this recruitment correlates with an increase in the recruitment and lifetime of CME coat protein AP-2 and late-arriving CME protein dynamin2. Furthermore, we find that CME sites that recruit Nef are more likely to recruit dynamin2, suggesting that Nef recruitment to CME sites promotes CME site maturation to ensure high efficiency in host protein downregulation.

## Introduction

Human immunodeficiency virus (HIV) depends on accessory viral proteins for its efficient replication and spread (Collins & Collins, 2014). HIV accessory proteins, including Nef, Vpu, Vif, Vpr, and Vpx, counteract host defenses by redirecting cellular pathways for the benefit of the virus (Collins & Collins, 2014; Malim & Bieniasz, 2012).

Here, we focus on the protein Nef, a non-enzymatic accessory protein encoded in the HIV genome (Allan et al., 1985). A major function of Nef is to downregulate host transmembrane proteins involved in antiviral restriction and immune response by modulating the trafficking routes of these membrane proteins (Pereira & daSilva, 2016; Buffalo, Iwamoto, et al., 2019). HIV-1 Nef downregulates specific host proteins, including CD3, CD4, CD28, MHC-I and SERINC5, by acting as an adaptor to physically bridge the target protein with the AP-1 and AP-2 adaptor complexes (Pereira & daSilva, 2016; Buffalo, Iwamoto, et al., 2019). The activities of Nef include internalizing surface proteins from the plasma membrane by clathrin-mediated endocytosis (CME), directing membrane proteins to lysosomal compartments for degradation, and sequestering membrane proteins at the Golgi (Pereira & daSilva, 2016; Buffalo, Iwamoto, et al., 2019). Nef function in the host cell is relatively well understood in a qualitative sense. While advances in live-cell imaging have made it possible to quantitatively follow CME with a high level of spatial and temporal precision (Cocucci et al., 2012; Mettlen et al., 2018; Dambournet et al., 2018; Pedersen et al., 2020), these powerful techniques have yet to be deployed to the study of Nef.

Studies described here focus on Nef’s interaction with the clathrin-mediated endocytosis (CME) pathway, the major pathway for plasma membrane protein internalization in mammalian cells (Bitsikas et al., 2014). Nef has been shown to downregulate the CD4 receptor (Guy et al., 1987; Inoue et al., 1993; Aiken et al., 1994) and the restriction factors SERINC3/5 (Usami et al., 2015; Rosa et al., 2015) and tetherin (in SIV but not HIV) (Zhang et al., 2009; Jia et al., 2009; Serra-Moreno et al., 2013), from the plasma membrane in a CME-dependent manner. AP-2, the major adaptor complex for CME (Jackson, COSB 2012; Traub & Bonifacino, CSHL 2013), is required for Nef downregulation of target proteins. Nef binds to AP-2 directly through a dileucine motif in a flexible disordered loop (Ren et al., 2014). In addition, structures of Nef complexed with AP-2 and the Nef-binding cytosolic tail of downregulated host factors have been solved (Buffalo, Sturzel, et al., 2019; Kwon et al., 2020).

In cells, colocalization of clathrin/AP-2, Nef, and Nef-downregulated host factors has been reported through live-cell and immunofluorescence microscopy, and was found to be dependent on a Nef dileucine motif (Greenberg et al., 1997; Greenberg et al., 1998; Burtey et al., 2007). Clathrin and Nef dynamics have been visualized in HeLa cells, where Nef was shown to be internalized from the plasma membrane together with clathrin (Burtey et al., 2007). These findings support a model in which Nef interacts with AP-2 to recruit its target proteins to CME sites for internalization from the plasma membrane. However, the quantitative details of how Nef affects CME dynamics, and how efficiently Nef recruits its target proteins to CME sites, have yet to be investigated in a living, physiologically relevant cell line. Such an analysis promises to provide novel insights into the mechanism of Nef action in CME.

In this study, we employ Jurkat cells, which are immortalized human T cells (Schwenk & Schneider, 1975; Abraham & Weiss, 2004), that were genome-edited to express fluorescent protein-tagged CME proteins AP-2 (*AP2M1*) and dynamin 2 (*DNM2*) at endogenous levels for faithful investigation of CME dynamics through live-cell microscopy. We selected the HIV-1 NA7 Nef patient isolate because it has been shown to potently promote CD4 internalization (Mariani & Skowronski, 1993; Mariani et al., 1996). Our quantitative analysis of CME dynamics revealed Nef-dependent changes in CME dynamics. Lastly, productivity of CME sites with or without Nef recruitment was assessed, revealing a Nef enhancement of the efficiency of target downregulation through the CME pathway.

## Results

### CME dynamics in genome-edited Jurkat cells

HIV Nef causes several different cell surface host factors to be removed from the cell surface by CME, but whether Nef affects CME dynamics is not known. To develop a cell line that would allow us to investigate whether HIV Nef affects CME dynamics, we genome-edited Jurkat cells, a T-cell leukemia cell line that resembles natural HIV hosts (Schwenk & Schneider, 1975; Abraham & Weiss, 2004). The resulting line expresses fluorescent protein-tagged endocytic proteins at endogenous levels. These cells were engineered to express the endocytic adapter protein AP-2 labeled with tagRFPt, and the late arriving scission factor dynamin 2 (DNM2) labeled with EGFP (Figure S1A and S1B) (Doyon et al., 2011; Hong et al., 2015). AP-2 is both the direct target of Nef and an established marker for clathrin-coat assembly (Boucrot et al., 2010; Cocucci et al., 2012). DNM2 serves as marker for vesicle scission (Grassart et al., 2014; Cocucci et al., 2014; Antonny et al., 2016; Merrifield et al., 2002) (Figure S1F).

Two imaging modalities were used to observe CME and Nef. The first was TIRF microscopy, which allows quantitative and sensitive imaging within ∼250 nm of the cell-substrate interface, facilitating the selective visualization of CME at a cell’s basal membrane (Merrifield et al., 2002). However, CME dynamics at the cell-substrate interface could be affected by factors such as substrate adhesion and membrane tension (Batchelder & Yarar, 2010). To measure CME at non-adhered surfaces, we also employed a complementary imaging method, HiLo microscopy (Tokunaga et al., 2008), to image the medial focal plane of cells, similar to what is done commonly with yeast (Kaksonen et al., 2003). We opted for HiLo microscopy over conventional confocal microscopy for this purpose due to the high signal-to-noise ratio and low photobleaching offered by the method (Tokunaga et al., 2008). Together, these two imaging modalities allowed us to quantitatively analyze CME dynamics in Jurkat cells. As in other cell types analyzed previously, AP-2 and DNM2 colocalized at cortical endocytic patches (Figure 1A and 1C).

**Figure 1.**
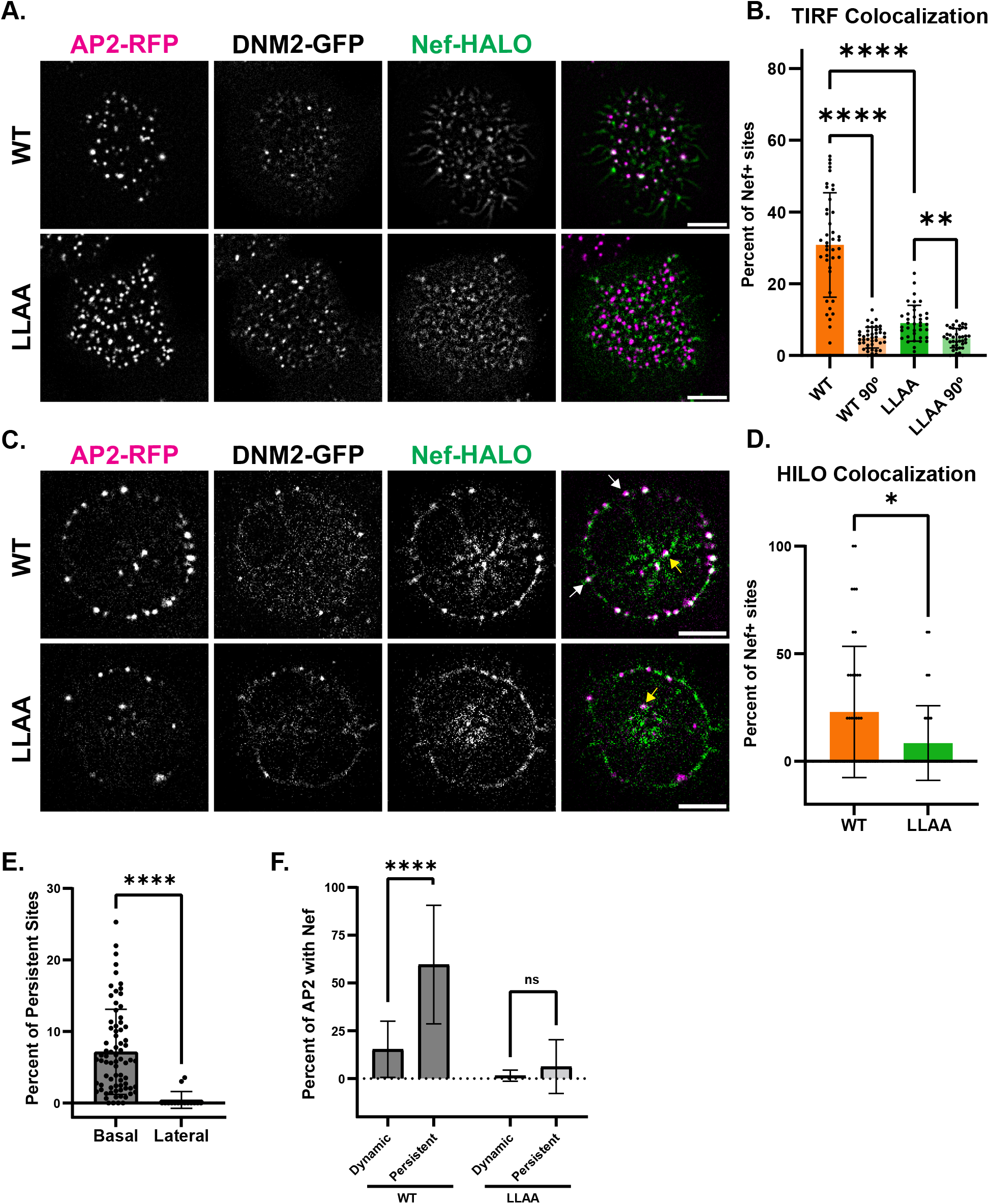
HIV Nef is recruited to CME sites. A. TIR-FM image of Jurkat cells endogenously expressing AP-2-RFP, DNM2-GFP, and transfected overnight with a construct expressing WT or LLAA Nef-HALO. Images were median filtered to highlight punctate structures. Scale bar is 5 μm. B. Quantification of the colocalization between AP-2 and WT/LLAA Nef observed in TIR-FM movie. WT/LLAA 90° conditions represent when reference images were rotated 90° to estimate random colocalization. The percentage of AP-2 pixels that overlap with Nef signal is plotted. AP-2 and WT Nef showed 31 ± 15% colocalization, while AP-2 and LLAA Nef showed 9 ± 5% colocalization. Each point represents a cell. **, **** indicates p value between 0.01 –0.001 and < 0.0001, respectively. C. Midfocal plane image of Jurkat cells endogenously expressing AP-2-RFP and DNM2-GFP, and transfected overnight with a construct expressing WT or LLAA Nef-HALO. Images were acquired through highly inclined and laminated optical sheet (HiLo) microscopy. Images were median filtered to highlight punctate structures. White arrows indicate cortical colocalization events. Yellow arrows indicate intracellular colocalization events. Scale bar is 5 μm. D. Quantification of the colocalization between AP-2 and WT/LLAA Nef observed in HiLo movie. Percentage of AP-2 puncta that recruit Nef was manually quantified and plotted. AP-2 and WT Nef showed 23 ± 30% colocalization while AP-2 and LLAA Nef showed 8 ± 17% colocalizaion. Each point represents a cell. * indicates p value between 0.01 – 0.05. E. Quantification of the percentage of persistent AP-2 events on the membrane regions in contact or not in contact with the cover glass. 7 ± 6% of AP-2 events were persistent on the portion of membrane attached to the cover glass and 0.4 ± 1% of AP-2 events were persistent on the portion of membrane not attached to the cover glass. Each point represents a cell. **** indicates p < 0.0001. F. Quantification of the colocalization of WT/LLAA Nef in dynamic and persistent AP-2 sites from TIR-FM movies. WT Nef showed 15 ± 15% colocalization with dynamic sites and 60 ± 31% colocalization with persistent sites. LLAA Nef showed 2 ± 3% colocalization with dynamic sites and 6 ± 14% colocalization with persistent sites. **** indicates p < 0.0001 and ns indicates p ≥ 0.05.

### Nef is specifically recruited to CME sites in Jurkat cells

Nef contains an AP-2-binding dileucine motif shown previously by both live cell imaging and fixed-cell immunostaining to be required for efficient recruitment to CME sites (Greenberg et al., 1997; Burtey et al., 2007). To test whether this motif is also required for Nef recruitment to CME sites in Jurkat cells, we transfected the genome-edited Jurkat cells with either WT Nef or the dileucine motif LLAA mutant Nef. To examine Nef recruitment at the membrane, Nef localization was assessed through both TIRF and HiLo microscopy (Figure 1A and 1C, Movie 2-5). TIRF microscopy results indicated that WT Nef shows 31 ± 15% colocalization with AP-2 while LLAA Nef shows only 9 +/-5% colocalization (Figure 1B). Random colocalization between Nef and AP-2 was estimated by rotating the AP-2 image 90 degrees and quantifying colocalization with its corresponding Nef image. Both WT and LLAA Nef show greater colocalization in non-rotated images compared to random colocalization in the rotated images (Figure 1B). For CME events on the portion of membrane not attached to the cover glass, WT Nef colocalized with AP-2 23 ± 30% of the time while LLAA Nef only colocalized with AP-2 8 ± 17% of the time (Figure 1D). The measured colocalization value for Nef and AP-2 was lower when assessed by HiLo microscopy on the portion of membrane not attached to the cover glass due to their lower fraction of persistent AP-2 events (Figure 1E). Persistent AP-2 events, lasting throughout the entire 5 minute movie, was mostly only present in TIRF movies and exhibited increased dileucine dependent colocalization with Nef at a much higher 60 ± 31% (Figure 1F). Both approaches confirm that Nef localizes to CME sites in a dileucine motif-dependent manner across all of the plasma membrane. Furthermore, these results suggest that Nef localization to CME sites is affected by contact between substrate and the plasma membrane, since increased colocalization between Nef and AP-2 was observed at persistent AP-2 events on the portion of membrane attached to the cover glass.

Unexpectedly, we also observed Nef and AP-2 colocalization at intracellular structures. These structures likely reflect at least two types of Nef-AP-2 interactions. Some such structures appear to be recently internalized clathrin-coated vesicles that have not been uncoated, since we were able to observe Nef and AP-2 internalize from plasma membrane together after DNM2 disassembly (Figure S2A). Intracellular structures exhibiting Nef-AP-2 colocalization were also observed in cells expressing LLAA Nef (Figure S2B). The nature of this second class of structure is presently unknown and may represent interactions between AP-2 and one of several organelles that recruit Nef (Pereira & daSilva, 2016).

### Nef recruitment increases AP-2 lifetime at CME sites

We classified CME events into three categories. The control category represents CME dynamics measured in untransfected AP-2-RFP DNM2-GFP Jurkat cells. The Nef+ and Nef-categories refer to CME sites in Nef expressing cells that do and do not recruit Nef, respectively. Since substrate adhesion may alter CME dynamics, we chose to first analyze the effect of Nef on CME dynamics via HiLo microscopy. To quantify CME dynamics at many CME sites, we generated kymographs around the circumference of these imaged through their medial focal planes (Figure 2A right). We quantified the lifetime that the fluorescent proteins resided on the cell periphery by measuring the length of each endocytic event on these kymographs (Figure 2A left). The lifetime analysis revealed that CME sites in untransfected cells or at sites of transfected cells that lack detectable Nef exhibited similar AP-2 lifetimes of 66 ± 21 and 64 ± 19 seconds, respectively, while Nef+ CME sites exhibited longer AP-2 lifetimes averaging 87 ± 23 seconds (Figure 2B). The LLAA Nef+ and Nef-CME sites displayed very similar lifetimes of 66 ± 28 and 65 ± 24 seconds, respectively, showing no statistically significant change in AP-2 lifetime correlated with Nef recruitment. These results show that recruitment of WT Nef to a CME site increases its lifetime.

**Figure 2.**
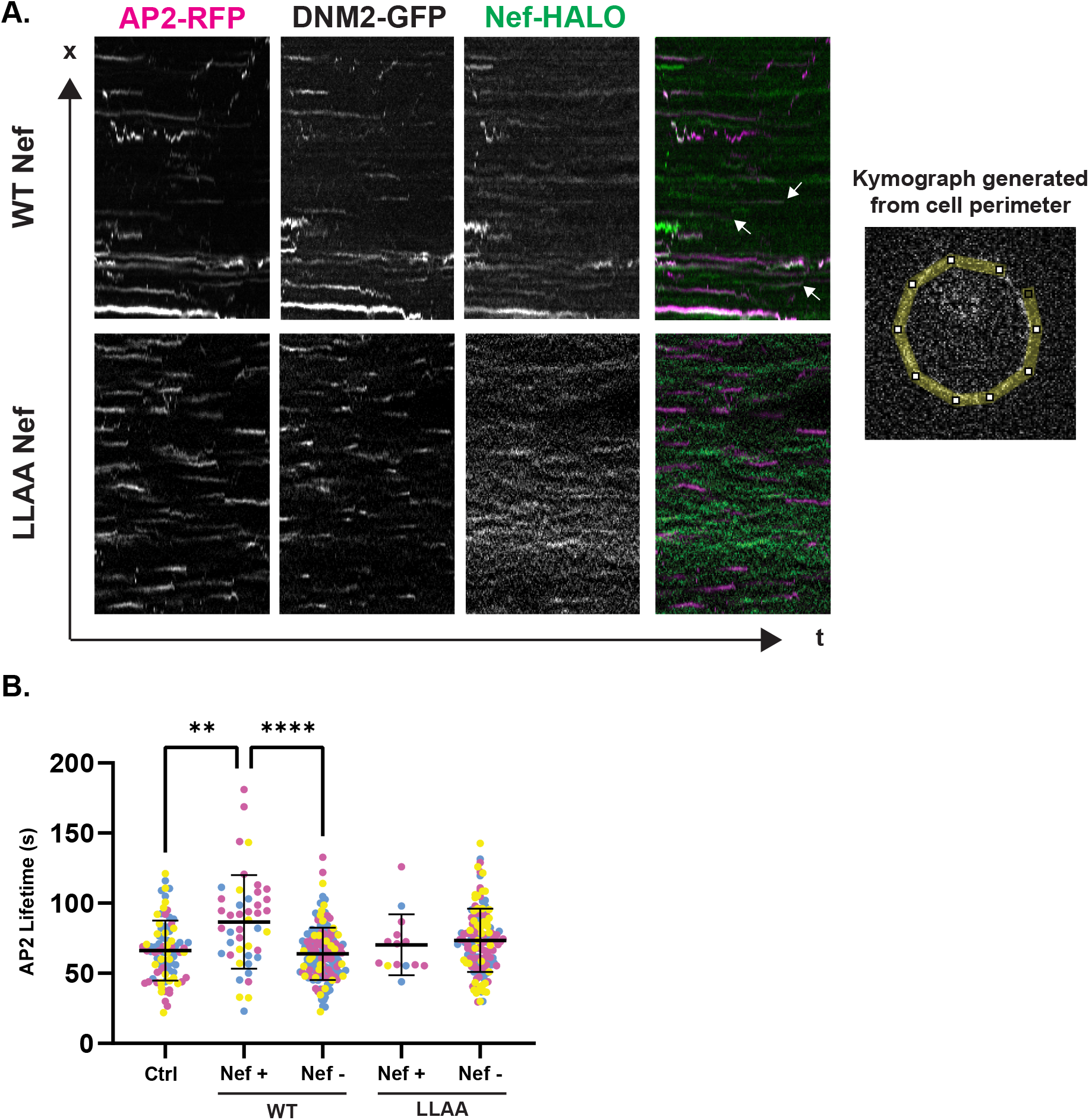
Nef positive CME sites display longer AP-2 lifetimes. A. Kymograph analysis of Jurkat cells endogenously expressing AP-2-RFP, DNM2-GFP and transfected overnight with WT or LLAA Nef-HALO. Each kymograph displays the circumference of a single cell in a 5-minute long HiLo microscopy movie. Nef positive CME events are annotated with white arrows. B. AP-2 lifetime quantification from kymograph analysis. The measured AP-2 lifetimes were 66 ± 21, 64 ± 19, 66 ± 28, and 65 ± 24 seconds, for control, WT Nef-, WT Nef+, LLAA Nef- and LLAA Nef+ events respectively. **, **** indicates p value 0.01 – 0.001 and <0.0001, respectively.

### Increased AP-2 and DNM2 recruitment and lifetimes observed at Nef-positive CME sites by TIRF microscopy

We turned to TIRF imaging for higher throughput lifetime and intensity analysis. We validated our analysis pipeline for CME events and Nef colocalization in TIRF movies by confirming that the analysis reported a similar amount of colocalization between WT Nef and AP-2 at dynamic CME sites as seen through HiLo microscopy, while exhibiting reduced colocalization between LLAA Nef and AP-2 (Figure S3A). We focused our analysis on dynamic events since persistent events were not often seen on the portion of membrane not attached to the cover glass. We determined that 15% of basal CME sites were Nef positive in WT Nef transfected cells, while only 1.6% were Nef positive in LLAA Nef transfected cells. The Nef positive events in LLAA showed reduced Nef lifetime and recruitment (Figure S3B and S3C). We further improved our approach by stably integrating an inducible form of HIV-Nef-HALO into the genome-engineered Jurkat cells to gain control over the duration and amount of Nef expression (Figure 3A, Figure S5, Movie 6).

**Figure 3.**
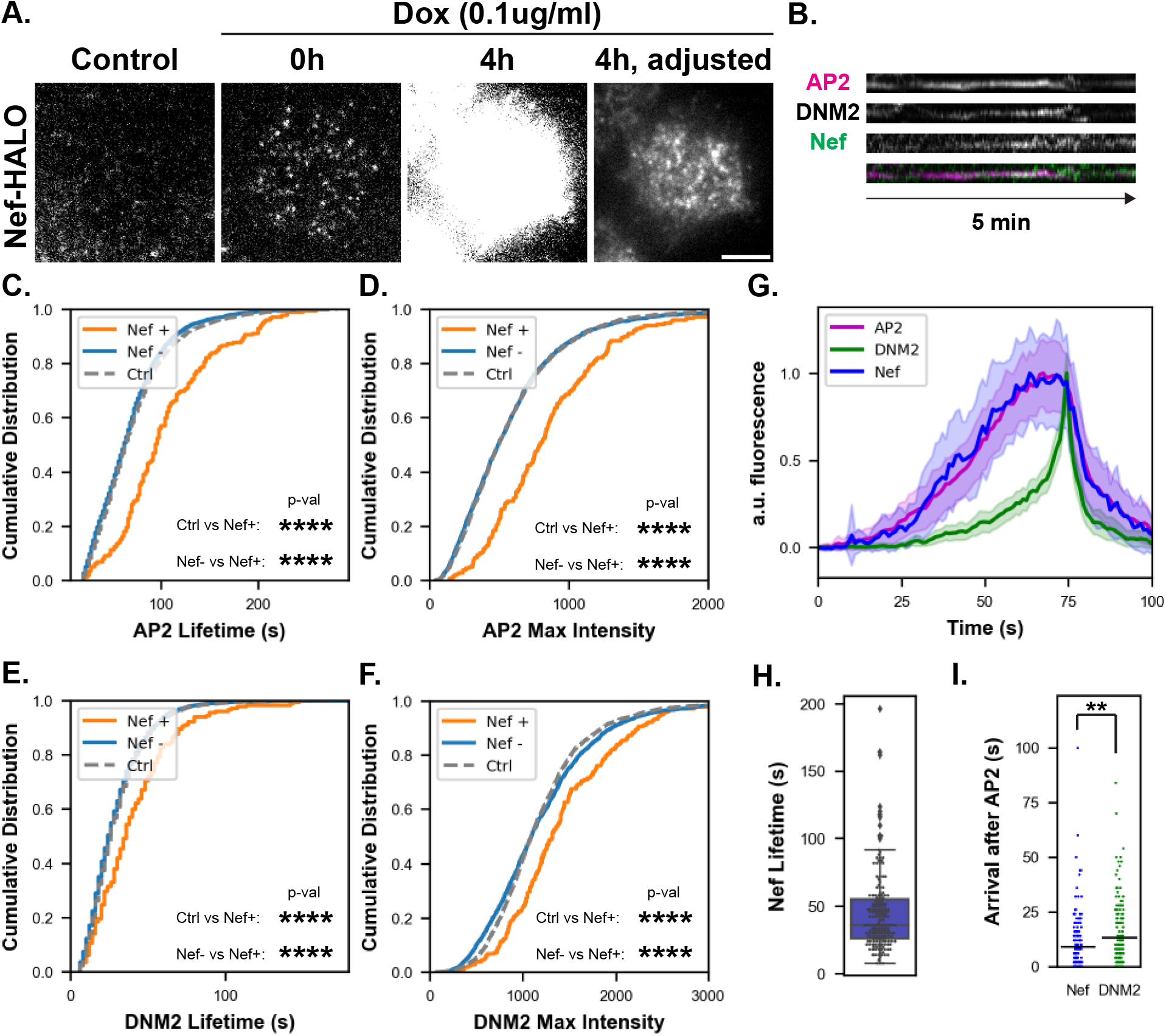
Nef positive CME sites display longer AP-2 and DNM2 lifetimes and higher maximum intensities. A. TIR-FM image of Jurkat cells endogenously expressing AP-2-RFP, DNM2-GFP, and inducibly expressing Nef-HALO. Control image shows background autofluorescence in the Nef imaging channel for parent AP-2-RFP, DNM2-GFP cell line. Adjusted image is the 4h induction Nef image with brightness adjusted to show punctate localization of Nef at the basal membrane. Scale bar is 5 μm. B. Kymograph from a 5 minute movie of a single Nef+ CME site in the AP-2-RFP, DNM2-GFP, Nef-HALO cell line after 4 hours of induction. C. Quantification of AP-2 lifetime as a cumulative distribution plot. The control (ctrl) CME events are those observed in the parental AP-2-RFP and DNM2-GFP Jurkat cell line. The Nef+ and Nef-CME events are those that recruit and do not recruit Nef in the inducible Nef Jurkat cell line after 4 hour induction. D. Quantification of AP-2 max intensity as a cumulative distribution plot. Same data set as C. E. Quantification of DNM2 lifetime as a cumulative distribution plot. Same data set as C. F. Quantification of DNM2 max intensity as a cumulative distribution plot. Same data set as C. For C-F, **** indicates p value <0.0001. G. Intensity over time of averaged Nef positive CME events with an AP-2 lifetime between 0-150 seconds. Outer lines of the curve represent standard deviations. Tracks were aligned to the maximum intensity of DNM2. H. Quantification of Nef lifetime calculated from Nef positive CME events 4h after induction. Nef lifetime was calculated to be 45 ± 30 seconds. Individual events are plotted as points. Outliers are plotted as diamonds. I. Quantification of time of first detection of Nef and DNM2 in Nef-positive events after the first AP-2 detection. Dataset consists of Nef positive events 4 hours after induction, regardless of DNM2 recruitment. Black bar indicates mean. ** indicates p value between 0.001 and 0.01.

We chose to analyze the effects of Nef on CME dynamics by inducing Nef expression for four hours, which was the earliest time point that showed marked increase in Nef expression levels by fluorescence microscopy compared to uninduced cells (Figure S4A and S4D). We first reproduced results from HiLo imaging that CME sites that recruit Nef exhibit increased AP-2 lifetimes (Figure 3C). Control CME sites had an AP-2 lifetime of 72 ± 40 seconds, Nef negative sites had an AP-2 lifetime of 68 ± 37 seconds, and Nef positive sites showed an increased AP-2 lifetime of 105 ± 51 seconds. Additionally, Nef positive CME sites showed increased AP-2 recruitment compared to Nef negative and control CME sites (control 586.7 ± 439.1 au, Nef-588.7 ± 470.4 au, Nef+ 877.2 ± 536.4 au) (Figure 3D). LLAA Nef+ sites showed little to no change compared to control or LLAA Nef-CME sites in AP-2 lifetime and recruitment (Figure S3D and S3E). Interestingly, similar increases in lifetime and recruitment at Nef+ sites were also observed for DNM2 (Figure 3E and 3F).

Next we determined the dynamics of Nef at CME sites (Figure 3B and 3G). Kymograph analysis from both HiLo and TIRF images showed that Nef tends to be recruited to CME sites shortly after AP-2, and persists until AP-2 disappears (Figure 2A and 3B). The cohort plot for CME also qualitatively matched the observations from the kymographs (Figure 3G). The AP-2 and DNM2 intensity profiles matched those of other cell lines (Hong et al., 2015; Dambournet et al., 2018). The intensity changes of Nef closely matched those of AP-2, further supporting the model that Nef affects CME through AP-2 interaction. Nef lifetime was measured to be 45 ± 30 seconds (Figure 3H). In summary, Nef+ sites show increased AP-2 and DNM2 recruitment and lifetime.

### Neither the number of Nef molecules at a CME site nor the total expression induction time predict AP-2 lifetime or amount

To gain further insight into how Nef affects CME dynamics, we asked whether the amount of Nef recruited to a CME site predicts the magnitude of the observed phenotypes. Using the Nef inducible cell line, Nef expression was induced for 0, 2, 4, 6, and 24h to vary the amount of Nef recruitment to CME sites. Nef expression increased with longer induction time (Figure S4A and S4B), as did the amount of Nef recruited to Nef positive sites and the percent of Nef positive sites (Figure S4C and S4D). However, there was no detectable difference in the magnitude of AP-2 lifetime or AP-2 amount recruited at different times of Nef induction (Figure S4E and S4F). These results suggest that the threshold for the full effect of Nef recruitment on CME site lifetime is low.

### Nef positive CME sites show increased productivity

Previous studies concluded that to varying degrees in different cell types, initiated CME events do not always proceed to productive vesicle formation. Characteristics of productive CME events include longer AP-2 lifetimes, higher AP-2 recruitment levels, and DNM2 recruitment in a sharp peak just before its disappearance (Loerke et al., 2009; Taylor et al., 2011; Hong et al., 2015; Aguet et al., 2013; Ehrlich et al., 2004). We sought to determine whether the effects of Nef on CME protein recruitment levels and lifetime might reflect increased productivity of Nef+ CME events.

One distinguishing feature of non-productive, or abortive, CME events is that their AP-2 lifetimes are often less than ∼20 seconds in length (Hong et al., 2015). Another is that CME events that are not completed fail to recruit DNM2 in a sharp peak prior to site disappearance. To this end, we first analyzed differences in AP-2 lifetimes of Nef+ and Nef-events through generation of histograms for all CME events, regardless of whether these events recruited DNM2 (Figure 4A). Strikingly, the histogram for Nef-events reveals predominantly short-lived AP-2 events, while the histogram for Nef+ events reveals a much larger fraction of CME events with longer AP-2 lifetimes. Such short-lived CME events have been attributed to abortive events that do not recruit late-arriving endocytic factors such as DNM2 and GAK/Auxilin, which mark vesicle scission and uncoating, respectively (Loerke et al., 2009; Aguet et al., 2013; Wang et al., 2020; He et al., 2020). Furthermore, control CME events showed a similar AP-2 lifetime distribution as Nef-events, while LLAA Nef+ events showed an intermediate AP-2 lifetime distribution (Figure 4B). Together, these observations indicate Nef+ events are much less likely to exhibit reduced AP-2 lifetimes, and that this effect is partially dependent on the interaction between Nef and AP-2.

**Figure 4.**
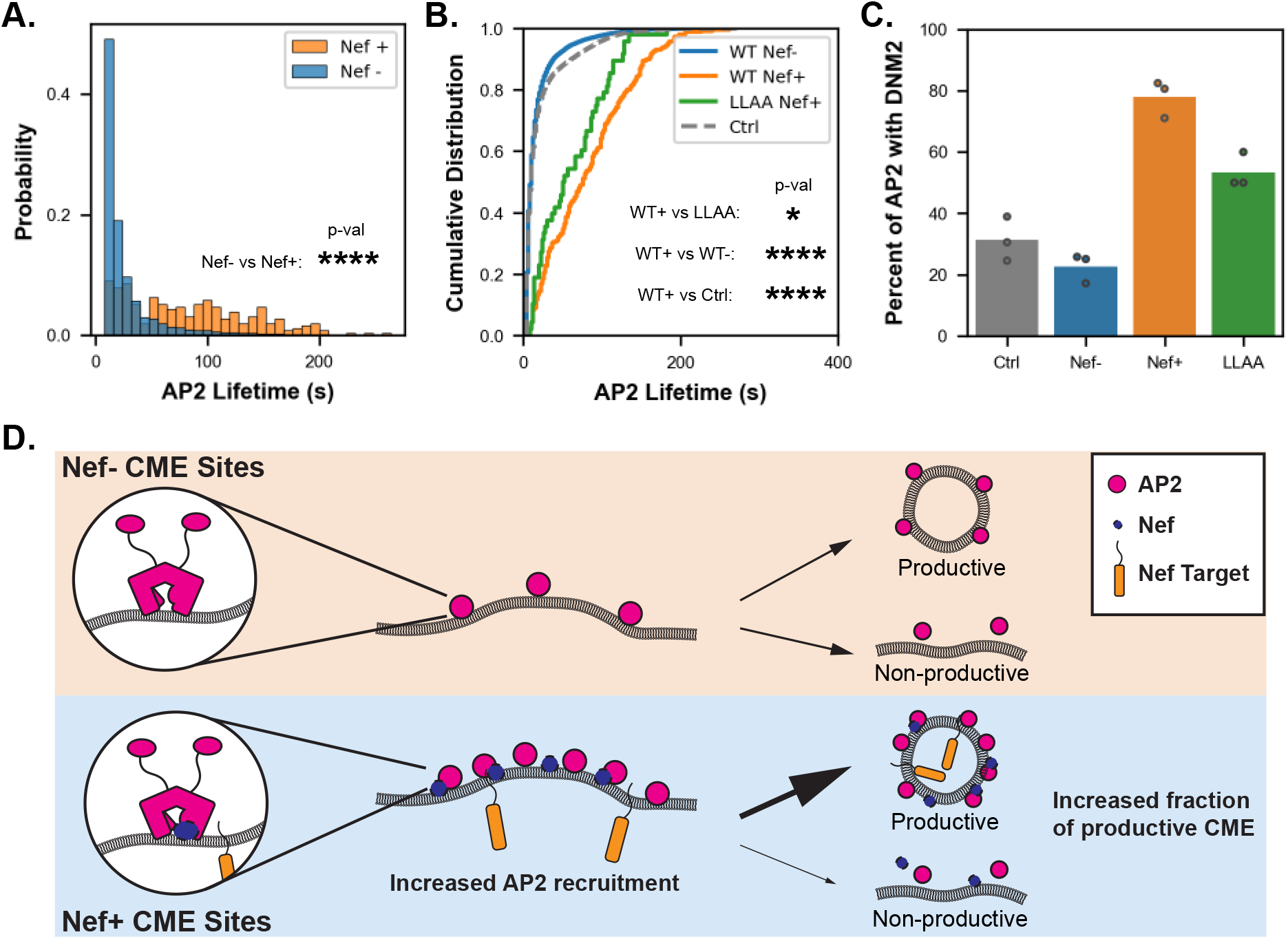
Nef recruitment increases DNM2-positive CME sites. A. Quantification of AP-2 lifetime as a histogram. Dataset consists of all Nef+ and Nef-CME events in WT Nef-transfected cells, regardless of DNM2 recruitment. **** indicates p value < 0.0001. B. Quantification of AP-2 lifetime as a cumulative distribution function. Dataset consists of all events in untransfected, WT Nef and LLAA Nef transfected cells, regardless of DNM2 recruitment. *, **** indicates p value <0.05 and < 0.0001, respectively. C. Bar plot showing the fraction of events at which DNM2 was recruited. Dataset consists of all events regardless of DNM2 recruitment from untransfected, WT Nef, or LLAA Nef transfected Jurkat cells. Points represent fraction of events that recruit DNM2 from separate biological replicates. D. Model for how Nef recruitment modifies CME kinetics. At Nef+ sites, Nef increases AP-2 recruitment. The Nef+ CME sites show increased productivity, meaning that more sites mature to vesicles with more AP-2 and presumably more cargo, resulting in more efficient internalization of Nef target proteins through CME.

If Nef is involved in determining CME site productivity, we reasoned that it should arrive at CME sites within the lifetime of abortive events. We determined that, on average, Nef is recruited around 18 seconds after AP-2 in Nef+ events. Since early recruitment of DNM2 has also been proposed to serve as a checkpoint in determining CME productivity, we measured the arrival of DNM2 in relation to AP-2 to further probe whether Nef is recruited at a sufficiently early time to influence CME productivity (Loerke et al., 2009). DNM2 was detected 26 seconds after AP-2 on average for Nef+ sites (Figure 3I). Thus, Nef is recruited during a time window when it could influence CME site productivity (Loerke et al., 2009; Aguet et al., 2013).

Finally, we assessed the productivity of Nef+ events by comparing the fraction of Nef+ events that recruited DNM2 to the Nef- and control events that recruited DNM2. DNM2 was recruited to 78% of Nef+ events and only 31% and 23% of control and Nef-events, respectively (Figure 4C). DNM2 was recruited to 53% of LLAA Nef+ events, showing an intermediate value that is consistent with the intermediate value of the AP-2 lifetime increase observed for LLAA Nef+ events (Figure 4C). This partial activity of LLAA Nef, along with the reduced but significant colocalization of LLAA Nef and AP-2 (Figure 1B), suggests that there might be additional factors of presently unknown identity that contribute to the interaction between Nef and CME machinery. In summary, these results suggest Nef+ CME sites are more productive than control or Nef-sites, and are consistent with the possibility that Nef arrives at CME sites early to increase productivity in an AP-2 dependent manner (Figure 4D).

## Discussion

Current literature supports a model in which HIV Nef downregulates CD4, SERINC5, and other membrane proteins from the cell surface by acting as an adaptor to connect AP-2 to the target protein. Here, we closely examined the kinetic aspect of this model in a cell line that resembles the natural biological host of HIV and expresses endogenously tagged fluorescent derivatives of CME proteins AP-2 and dynamin2.

Nef localized to CME sites in a dileucine motif-dependent manner in the genome-edited Jurkat cells, consistent with previous findings. We noticed that in the region of the plasma membrane that was not attached to the cover glass almost all CME sites were dynamic, appearing and disappearing. However, 7% of CME events on regions of plasma membrane not attached to the cover glass persisted for the entirety of our 5-minute movies. These persistent sites showed greatly increased Nef recruitment compared to dynamic sites. Together, these observations indicate that Nef recruitment to CME sites may be sensitive to contact between the infected-host cell and the extracellular environment. Additionally, we also observed colocalization between Nef and AP-2 away from the plasma membrane, in the cell interior. Some of these structures appear to be newly released endocytic vesicles that have not yet been uncoated. However, colocalization was also observed to a similar degree when the Nef dileucine motif was mutated, suggesting that at least those colocalization events were not recently released clathrin-coated vesicles. Such interactions between clathrin-coated vesicles and Nef positive endosomal organelles might therefore represent a previously unappreciated site at which Nef can modulate cellular traffic.

AP-2 and dynamin2 lifetimes were greater at Nef positive CME sites compared to Nef negative sites in the same cells. Therefore, the increased lifetime appears to be a direct consequence of Nef presence at a CME site rather than to changes in the overall cellular environment. The upper limit of observed AP-2 recruitment and lifetime was comparable across control, Nef-, and Nef+ sites, suggesting that the magnitude of change in these parameters of Nef+ CME sites could be restricted to a pre-existing range within the control CME population (Figure S4E and S4F).

While the upper end of the Nef effect on AP-2 and dynamin2 recruitment and dynamics can be explained by constraints on the CME site, the lack of correlation between the amount of Nef recruited and AP-2 recruitment/lifetime was unexpected. An intriguing possibility is that there is a local amplification of Nef effects at the CME site such that only a small amount of Nef is necessary to saturate the effect. Outside of its trafficking functions, Nef has been shown to modulate signaling through interaction with various kinases, such as activating Src family kinases through binding of Nef’s proline-rich domain to the negative regulatory SH3 domains of Src family kinases (Saksela et al., 1995). On the other hand, Nef mutations in the proline rich domain involved in binding the SH3 domains of Src family kinases do not interfere with downregulation of CD4 through CME (Saksela et al., 1995). Many CME proteins such as dynamin2 also interact through a multivalent SH3 and proline-rich domain network at the CME site, raising the possibility that the observed Nef-dependent changes might involve this network as well (Li et al., 2012; Sun et al., 2017). A third explanation for the lack of correlation between Nef levels and effects is the possibility that HIV Nef functions in a target sensitive manner. The closely related SIV Nef has been shown to form a tripartite complex consisting of the AP-2 complex, the cytoplasmic tail of the target host protein, and itself. Whether the interaction between Nef and AP-2 alone is sufficient to effect CME dynamics is not clear (Buffalo, Sturzel, et al., 2019; Kwon et al., 2020). A mixed population of target-bound Nef and target-free Nef can explain the lack of strong correlation between Nef’s levels and the magnitude of its effect on CME.

Finally, we observed that Nef increases CME site productivity. Non-productive or abortive events are characterized by low accumulation of endocytic coat proteins and lack of late CME protein recruitment, such as the final burst of dynamin2 recruitment and subsequent arrival of uncoating factors (Ehrlich et al., 2004; Aguet et al., 2013; Hong et al., 2015; He et al., 2020). Here, we showed that Nef-positive CME sites have significantly increased AP-2 lifetimes and recruitment levels, and are around 50% more likely to recruit detectable dynamin2 in a late peak. The majority of Nef is recruited to CME sites within 18 seconds of AP-2 recruitment, meaning that Nef is present within the lifetime of both abortive and productive events, the time window when it could influence CME site productivity. WT Nef-positive CME sites recruited more dynamin2 than LLAA Nef-positive CME sites, suggesting that the dileucine motif interaction between Nef and AP-2 plays a role in regulating CME productivity. Importantly, the dileucine motif interaction between Nef and AP-2 is similar to how some membrane receptors interact with AP-2 to be internalized (Ren et al., 2014; Kelly et al., 2008). Previous studies have demonstrated that overexpression or increase in local concentration of membrane proteins harboring the tyrosine motif, a different AP-2 binding motif, can increase CME productivity (Loerke et al., 2009; Liu et al., 2010; Kadlecova et al., 2017). An analogous mechanism where the Nef dileucine motif mimics membrane protein binding with AP-2 might increase CME productivity. Intriguingly, it was reported that overexpression of tyrosine motif-containing membrane proteins can increase the nucleation of CME events while overexpression of dileucine motif harboring membrane proteins cannot (Kadlecova et al., 2017). HIV Nef appears to have lost a tyrosine motif that is present in SIV Nef, and lacks the ability to downregulate CD3 from the cell surface via CME (Manrique et al., 2017). The differences in how AP-2 binding motifs influence CME dynamics and Nef target protein internalization may have played a role in Nef evolution.

In conclusion, we report that Nef increases the recruitment and lifetimes of CME proteins AP-2 and dynamin2 to CME sites. The effects of Nef on the AP-2 protein recruitment and lifetime were dependent on the dileucine motif of Nef. Finally, we observed that Nef-positive CME sites exhibit increased productivity. Given these observations, we propose that Nef promotes membrane protein downregulation at least in part by increasing CME site productivity through AP-2 stabilization.

## Methods

### Cloning

The donor vector for endogenously tagging the *dynamin2* gene was generated by fusing the pCR8 backbone, 5’ DNM2 homology arm, GTSGGS linker, tagGFP2, and 3’ DNM2 homology arm via Gibson assembly. The donor vector for endogenously tagging the *AP2M1* gene was generated by fusing the pCR8 backbone, 5’ AP-2 homology arm, GTSGGS linker, tagRFPt, GMDELYKASGSGT linker, and 3’ AP-2 homology arm via Gibson assembly. Homology arms of genes were obtained from genomic PCR. NA7-Nef-HALO was generated by fusing the pcDNA5 backbone with the codon-optimized coding sequence of the NA7 Nef (protein ID: ABB51086.1), SGGTG linker, and HALO tag via Gibson assembly. The pcDNA5 NA7-Nef-HALO LLAA construct was generated by mutating L164 and L165 into alanines via site-directed mutagenesis from the pcDNA5 NA7-Nef-HALO construct. The pInducer20 NA7-Nef- HALO was generated by cloning the NA7-Nef-HALO insert into the pENTR1A vector, followed by gateway assembly of the insert into the pInducer20 packaging vector (Meerbrey et al. 2011).

### Cell culture

Experiments were conducted using Jurkat E6-1 cells, originally obtained from the UCB Cell Culture Facility. The cells were cultured in RPMI-1640 Media supplemented with 10% FBS, Pen-Strep, and Sodium Pyruvate and grown at 37°C and 5% CO2. Cells were maintained at densities below 2e6 cells/ml during passaging and experiments were conducted within 20 passages from the generation of the cell line. Stable cell lines inducibly expressing Nef were selected with G418 concentration of 800μg/ml and maintained with concentration of 400μg/ml. MDA-MB-231 cells were cultured in DMEM F12 Glutamax supplemented with 10% FBS and Pen-Strep. For lentiviral production, HEK293T cells were cultured in DMEM/F-12 media supplemented with 10% FBS, Pen-Strep, and non-essential amino acids on 10cm plates.

Jurkat cells were obtained from the Berkeley Cell Culture Facility. STR profiling of AP2-RFP, DNM2-GFP and AP2-RFP, DNM2-GFP, NA7-HALO Jurkat cell lines was conducted for cell line authentication. Cell lines were tested for mycoplasma before use in microscopy experiments.

### Electroporation

For genome editing, Jurkat cells were electroporated using the Amaxa Nucleofector 2b, protocol X-01. For transient transfections, Jurkat cells were electroporated using the Ingenio electroporation solution following the manufacturer’s instructions (Mirus 50111).

Electroporation was conducted using the Biorad Gene Pulser Xcell System using 0.4cm cuvettes. 1e6 – 2e6 Jurkat cells were transfected per condition. The settings for the electroporation used were voltage of 180, pulse length of 10ms, and single pulse.

### Genome editing

Jurkat cells were genome-edited by electroporation of the Cas9-crRNAtracrRNA complex using the Amaxa Nucleofector 2b electroporator. Donor vectors for genome-editing have homology arms ranging from 700-1000 bp, and were electroporated with the Cas9 complex. The gRNA sequence TTGCTTCCCGCTGCAAGCAG was used to target *AP2M1*, and the gRNA sequence CCTGCTCGACTAGGCCTCGA was used to target *DNM2.* Jurkat cells were cultured for 3 days after electroporation and sorted into 96-well plates for clonal expansion using fluorescent signal to isolate edited clones using the BD Bioscience Influx sorter (BD Bioscience). Both alleles of AP-2 and one allele of DNM2 were tagged in the final cell line.

The *S. Pyogenes* NLS-Cas9 was purified in the University of California Berkeley QB3 Macrolab. Sorting was conducted in the University of California Berkeley CRL flow cytometry facility.

### Lentiviral production and infection

The pInducer20-NA7-Nef-Halo vector was packaged into lentiviruses in HEK293T cells by co-transfection with 2^nd^ generation lentiviral packaging plasmids (Addgene 182993 and 244397) using TransIT-LT1 transfection reagent according to the manufacturer’s instructions (Mirus 2304). Viral supernatants were collected for 3 days and filtered through a 0.45μm PES filter. The collected supernatant was concentrated with LentiX concentrator following the manufacturer’s instructions (Takara 631232). The virus containing pellet was then resuspended in 1ml of the supernatant and stored in 100μl aliquots at -80°C.

To generate stable Jurkat cell lines, Jurkat cells were sub-cultured in a well of a 6-well plate. An aliquot of the concentrated virus was directly pipetted into the well and cells were incubated with virus overnight. The following day, the media was replaced with media containing 800μg/ml G418 for selection. The infected Jurkat cells were selected in G418 for 2 weeks before use in experiments. After 2 weeks, the stable Jurkat cell lines were cultured in 400μg/ml G418.

### Live-cell microscopy

Jurkat cells were incubated with 100nM JF635 dye at least 1 hour prior to imaging if expressing or being compared to cell lines expressing HALO-tagged proteins. NA7 Nef expression was induced by treating cells with 0.1μg/ml Doxycycline for the desired duration. AP-2-RFP DNM2- GFP and the uninduced AP-2-RFP DNM2-GFP NA7-Nef-HALO Jurkat cell lines were incubated with an equal volume of DMSO instead for 24 hours.

All imaging was conducted on 8 well chambered glass slides (Cellvis C8-1.5H-N), coated with 1mg/ml Poly-L-Lysine. Plates were coated for 10 min and washed 3 times with PBS. Jurkat cells were transferred to imaging media (growth media without phenol red) and allowed to adhere on the well for 10 minutes prior to imaging.

Images were acquired through total internal reflection fluorescence (TIRF) microscopy using a Nikon Ecliplse Ti2 inverted microscope. The TIRF scope was equipped with a LUNF 4-line laser launch (Nikon Instruments) and an iLas2 RING-TIRF module (Gataca Systems). The CFI60 60x Apo TIRF objective and a Hamamatsu Orca Fusion Gen III sCMOS camera were used to capture images. All movies were acquired at 37°C sequentially over 5 min at 2 second intervals and 100ms exposure for each channel. The system was controlled with NIS-Elements software.

For HiLo microscopy, Jurkat cells were prepared as described and the excitation laser was angled at 55 degrees. The plane chosen for imaging was the focal plane closest to the bottom of the coverslip where fluorescence from the bottom surface of the cell could not be observed. The incubation condition and other imaging parameters were the same as in TIRF.

All datasets consist of three movies per condition per day, taken on three separate days. For Hilo imaging, at least 30 cells and 85 CME events were analyzed for each condition. For TIRF imaging, at least 39 cells were analyzed for each condition in transfection experiments. 2147, 1074, and 1356 CME events were analyzed in control, WT-NA7 expressing, and LLAA-NA7-Nef expressing Jurkat cells, respectively. At least 141 cells were analyzed for each condition in induction experiments. 3010, 4577, 3332, 2684, 2611, and 2510 CME events were analyzed in control, 0h, 2h, 4h, 6h, and 24h Nef induced Jurkat cells.

### Cell lysis and Western blot

1e6 – 2e6 Jurkat cells were washed with PBS and resuspended in 24μl of lysis buffer, consisting of 50 mM Hepes pH 7.4, 150mM NaCl, 1mM MgCl_2_, 1% NP40, phosSTOP (Roche 4906845001), and protease inhibitor (Roche 1183617001). Cells were lysed for 15 minutes on ice with gentle agitation every 5 minutes. Lysed cells were then centrifuged at 4°C and 13,000rpm for 15 minutes. 18 μl of the supernatant was collected as lysate. The lysate was mixed with Laemmeli reducing buffer with 5% β-mercaptoethanol. 10 μl of the final solution was loaded onto a 10% acrylamide gel for SDS-PAGE analysis. Proteins resolved on the SDS-PAGE gel was transferred onto a nitrocellulose membrane for immunoblotting via wet transfer. Blots were blocked with 5% milk in tris-buffered saline (TBS) and incubated with primary antibody. The antibodies used for western blotting were: anti-HALO (1:1000 in 0.5% milk TBST, 4°C o/n, Promega G9211), anti-GAPDH (1:10000 in TBST, RT 1 hr, ab9485), anti-AP-2M1 (1:500 in 0.5% milk TBST, 4°C o/n, Abcam, ab75995), and anti-Tag(CGY)FP (1:2000 in 0.5% milk TBST, 4°C o/n, Evrogen AB121). Blots were then incubated with IRDye 800/680 conjugated antibodies (1:10000 in 5% milk TBST, RT 1hr, 926-32212 926-68071) and imaged on the LICOR Odyssey scanner.

### Image Processing

Prior to kymograph and colocalization analysis through imageJ, images were background subtracted (subtract mean value of box outside cell from all pixels), corrected for photobleaching if applicable (Fiji bleach correction, exponential fitting method), and cytoplasmic background subtracted (Median filter, r = 6px, was subtracted from image) to clearly visualize punctate structures.

### Kymograph Analysis of endocytic events

To create kymographs of mid-focal plane imaging movies an average intensity Z-projection of these movies was generated and a drawing a 20 pixel wide line around circumference of cells. Intensity change of the movie within the line was projected across time. Lifetime measurements were made by drawing a horizontal line over the observed CME events in imageJ and measuring its length. The beginning and end of events were determined by when intensity levels were at least 3x background levels. Events were categorized as persistent if the AP-2 signal was present throughout the entire 5 minute movie. Events were categorized as dynamic if the AP-2 signal appeared and disappeared within the 5 minute movie.

### Colocalization Analysis

Colocalization was evaluated using three different methods. For HiLo images, colocalization was manually assessed through kymograph analysis. For TIRF images, colocalization analysis for all events was conducted through imageJ. Individual cells were segmented into square regions and separately assessed for colocalization. After processing, average intensity projections were generated for each channel. Masks for punctate structures were generated using the imageJ “Find Maxima” function (prominence setting of 10, with tolerance). The pixel overlap of the resulting masks was assessed for pairs of channels to test for colocalization. As a negative control, the mask generated from the AP-2 coat channel was rotated 90 degrees and compared to masks from other channels, establishing a baseline for chance colocalization.

### Particle tracking analysis

For particle tracking analysis, cells were cropped individually and separated into cell specific folders for analysis using the cmeAnalysis MATLAB package. For tracking, gaussian PSF model fitting detection with a gap length of 2, tracking minimum radius of 3, and tracking maximum radius of 6. Tracks that showed an AP-2 lifetime greater than 20 seconds, mean squared displacement <0.02μm^2^, and at least 3 consecutive significant (pval < 0.01) detections of DNM2 were considered valid tracks. Tracks that showed at least 4 consecutive significant (pval < 1e-25) detections of Nef that averaged within < 1.25 pixels away from AP-2 were considered to be Nef positive tracks. These parameters for assessing Nef colocalization were determined by manually examining montages of detected Nef colocalization events, and by comparing the number of detected Nef positive events in WT Nef and LLAA Nef transfected samples. Dynamic events and persistent events were filtered by analyzing category 1 and 4 tracks in the CME analysis software, respectively. Lifetimes of DNM2 and Nef were calculated with a tolerable gap length of 2. For figure 4, tracks were considered to be DNM2 positive if there were at least 3 consecutive significant detections of DNM2 as well as a DNM2 maximum intensity higher than the 75th percentile of CME events that are shorter than 20 seconds.

### Scripts used for imaging analysis (Code Availability)

All user-made scripts for imageJ, MATLAB, and python can be accessed here: https://github.com/yuichiro-iwamoto/Nef-CME-analysis

### Statistical analysis

Statistical analysis was performed by using the python package sciPy or with Prism 9. Non-parametric statistical tests were used to evaluate significance so no assumption about data distributions were made. Specifically, Mann Whitney U test was used to evaluate significance of two groups and the Kruskal-Wallis test was used to evaluate significance when multiple comparisons were made. Data were visualized using the python package seaborn and Prism 9. All reported values are indicated as mean ± standard deviation.

## Acknowledgments

We thank Xuefeng Ren for generating the NA7 Nef construct, Frank Kirchhoff for helpful suggestions and Jonathan Wong for critical reading of the manuscript. This research was supported by NIH grants R01 AI120691 (J.H.H.), P50 AI150476 (D.G.D. and J.H.H.), National Institute of General Medical Sciences (NIGMS) grant R35 GM118149 to D.G.D.

## References

Abraham, R. T., & Weiss, A. (2004). Jurkat T cells and development of the T-cell receptor signalling paradigm. Nat Rev Immunol, 4(4), 301–308. https://doi.org/10.1038/nri1330

Aguet, F., Antonescu, C. N., Mettlen, M., Schmid, S. L., & Danuser, G. (2013). Advances in analysis of low signal-to-noise images link dynamin and AP2 to the functions of an endocytic checkpoint. Dev Cell, 26(3), 279–291. https://doi.org/10.1016/j.devcel.2013.06.019

Aiken, C., Konner, J., Landau, N. R., Lenburg, M. E., & Trono, D. (1994). Nef induces CD4 endocytosis: requirement for a critical dileucine motif in the membrane-proximal CD4 cytoplasmic domain. Cell, 76(5), 853–864. https://doi.org/10.1016/0092-8674(94)90360-3

Allan, J. S., Coligan, J. E., Lee, T. H., McLane, M. F., Kanki, P. J., Groopman, J. E., & Essex, M. (1985). A new HTLV-III/LAV encoded antigen detected by antibodies from AIDS patients. Science, 230(4727), 810–813. https://doi.org/10.1126/science.2997921

Antonny, B., Burd, C., De Camilli, P., Chen, E., Daumke, O., Faelber, K., Ford, M., Frolov, V. A., Frost, A., Hinshaw, J. E., Kirchhausen, T., Kozlov, M. M., Lenz, M., Low, H. H., McMahon, H., Merrifield, C., Pollard, T. D., Robinson, P. J., Roux, A., & Schmid, S. (2016). Membrane fission by dynamin: what we know and what we need to know. EMBO J, 35(21), 2270–2284. https://doi.org/10.15252/embj.201694613

Batchelder, E. M., & Yarar, D. (2010). Differential requirements for clathrin-dependent endocytosis at sites of cell-substrate adhesion. Mol Biol Cell, 21(17), 3070–3079. https://doi.org/E09-12-1044 [pii] 10.1091/mbc.E09-12-1044

Bitsikas, V., Correa, I. R., Jr., & Nichols, B. J. (2014). Clathrin-independent pathways do not contribute significantly to endocytic flux. Elife, 3, e03970. https://doi.org/10.7554/eLife.03970

Boucrot, E., Saffarian, S., Zhang, R., & Kirchhausen, T. (2010). Roles of AP-2 in clathrin-mediated endocytosis. PLoS One, 5(5), e10597. https://doi.org/10.1371/journal.pone.0010597

Buffalo, C. Z., Iwamoto, Y., Hurley, J. H., & Ren, X. (2019). How HIV Nef Proteins Hijack Membrane Traffic To Promote Infection. J Virol, 93(24). https://doi.org/10.1128/JVI.01322-19

Buffalo, C. Z., Sturzel, C. M., Heusinger, E., Kmiec, D., Kirchhoff, F., Hurley, J. H., & Ren, X. (2019). Structural Basis for Tetherin Antagonism as a Barrier to Zoonotic Lentiviral Transmission. Cell Host Microbe, 26(3), 359–368 e358. https://doi.org/10.1016/j.chom.2019.08.002

Burtey, A., Rappoport, J. Z., Bouchet, J., Basmaciogullari, S., Guatelli, J., Simon, S. M., Benichou, S., & Benmerah, A. (2007). Dynamic interaction of HIV-1 Nef with the clathrin-mediated endocytic pathway at the plasma membrane. Traffic, 8(1), 61–76. https://doi.org/10.1111/j.1600-0854.2006.00512.x

Cocucci, E., Aguet, F., Boulant, S., & Kirchhausen, T. (2012). The first five seconds in the life of a clathrin-coated pit. Cell, 150(3), 495–507. https://doi.org/10.1016/j.cell.2012.05.047

Cocucci, E., Gaudin, R., & Kirchhausen, T. (2014). Dynamin recruitment and membrane scission at the neck of a clathrin-coated pit. Mol Biol Cell, 25(22), 3595–3609. https://doi.org/10.1091/mbc.E14-07-1240

Collins, D. R., & Collins, K. L. (2014). HIV-1 accessory proteins adapt cellular adaptors to facilitate immune evasion. PLoS Pathog, 10(1), e1003851. https://doi.org/10.1371/journal.ppat.1003851

Dambournet, D., Sochacki, K. A., Cheng, A. T., Akamatsu, M., Taraska, J. W., Hockemeyer, D., & Drubin, D. G. (2018). Genome-edited human stem cells expressing fluorescently labeled endocytic markers allow quantitative analysis of clathrin-mediated endocytosis during differentiation. J Cell Biol, 217(9), 3301–3311. https://doi.org/10.1083/jcb.201710084

Doyon, J. B., Zeitler, B., Cheng, J., Cheng, A. T., Cherone, J. M., Santiago, Y., Lee, A. H., Vo, T. D., Doyon, Y., Miller, J. C., Paschon, D. E., Zhang, L., Rebar, E. J., Gregory, P. D., Urnov, F. D., & Drubin, D. G. (2011). Rapid and efficient clathrin-mediated endocytosis revealed in genome-edited mammalian cells. Nat Cell Biol, 13(3), 331–337. https://doi.org/10.1038/ncb2175

Ehrlich, M., Boll, W., Van Oijen, A., Hariharan, R., Chandran, K., Nibert, M. L., & Kirchhausen, T. (2004). Endocytosis by random initiation and stabilization of clathrin-coated pits. Cell, 118(5), 591–605. https://doi.org/10.1016/j.cell.2004.08.017

Grassart, A., Cheng, A. T., Hong, S. H., Zhang, F., Zenzer, N., Feng, Y., Briner, D. M., Davis, G. D., Malkov, D., & Drubin, D. G. (2014). Actin and dynamin2 dynamics and interplay during clathrin-mediated endocytosis. J Cell Biol, 205(5), 721–735. https://doi.org/10.1083/jcb.201403041

Greenberg, M., DeTulleo, L., Rapoport, I., Skowronski, J., & Kirchhausen, T. (1998). A dileucine motif in HIV-1 Nef is essential for sorting into clathrin-coated pits and for downregulation of CD4. Curr Biol, 8(22), 1239–1242. https://doi.org/10.1016/s0960-9822(07)00518-0

Greenberg, M. E., Bronson, S., Lock, M., Neumann, M., Pavlakis, G. N., & Skowronski, J. (1997). Co-localization of HIV-1 Nef with the AP-2 adaptor protein complex correlates with Nef-induced CD4 down-regulation. EMBO J, 16(23), 6964–6976. https://doi.org/10.1093/emboj/16.23.6964

Guy, B., Kieny, M. P., Riviere, Y., Le Peuch, C., Dott, K., Girard, M., Montagnier, L., & Lecocq, J. P. (1987). HIV F/3’ orf encodes a phosphorylated GTP-binding protein resembling an oncogene product. Nature, 330(6145), 266–269. https://doi.org/10.1038/330266a0

He, K., Song, E., Upadhyayula, S., Dang, S., Gaudin, R., Skillern, W., Bu, K., Capraro, B. R., Rapoport, I., Kusters, I., Ma, M., & Kirchhausen, T. (2020). Dynamics of Auxilin 1 and GAK in clathrin-mediated traffic. J Cell Biol, 219(3). https://doi.org/10.1083/jcb.201908142

Hong, S. H., Cortesio, C. L., & Drubin, D. G. (2015). Machine-Learning-Based Analysis in Genome-Edited Cells Reveals the Efficiency of Clathrin-Mediated Endocytosis. Cell Rep, 12(12), 2121–2130. https://doi.org/10.1016/j.celrep.2015.08.048

Inoue, M., Koga, Y., Djordjijevic, D., Fukuma, T., Reddy, E. P., Yokoyama, M. M., & Sagawa, K. (1993). Down-regulation of CD4 molecules by the expression of Nef: a quantitative analysis of CD4 antigens on the cell surfaces. Int Immunol, 5(9), 1067–1073. https://doi.org/10.1093/intimm/5.9.1067

Jia, B., Serra-Moreno, R., Neidermyer, W., Rahmberg, A., Mackey, J., Fofana, I. B., Johnson, W. E., Westmoreland, S., & Evans, D. T. (2009). Species-specific activity of SIV Nef and HIV-1 Vpu in overcoming restriction by tetherin/BST2. PLoS Pathog, 5(5), e1000429. https://doi.org/10.1371/journal.ppat.1000429

Kadlecova, Z., Spielman, S. J., Loerke, D., Mohanakrishnan, A., Reed, D. K., & Schmid, S. L. (2017). Regulation of clathrin-mediated endocytosis by hierarchical allosteric activation of AP2. J Cell Biol, 216(1), 167–179. https://doi.org/10.1083/jcb.201608071

Kaksonen, M., Sun, Y., & Drubin, D. G. (2003). A pathway for association of receptors, adaptors, and actin during endocytic internalization. Cell, 115(4), 475–487. https://doi.org/10.1016/s0092-8674(03)00883-3

Kelly, B. T., McCoy, A. J., Spate, K., Miller, S. E., Evans, P. R., Honing, S., & Owen, D. J. (2008). A structural explanation for the binding of endocytic dileucine motifs by the AP2 complex. Nature, 456(7224), 976–979. https://doi.org/10.1038/nature07422

Kwon, Y., Kaake, R. M., Echeverria, I., Suarez, M., Karimian Shamsabadi, M., Stoneham, C., Ramirez, P. W., Kress, J., Singh, R., Sali, A., Krogan, N., Guatelli, J., & Jia, X. (2020). Structural basis of CD4 downregulation by HIV-1 Nef. Nat Struct Mol Biol, 27(9), 822–828. https://doi.org/10.1038/s41594-020-0463-z

Li, P., Banjade, S., Cheng, H. C., Kim, S., Chen, B., Guo, L., Llaguno, M., Hollingsworth, J. V., King, D. S., Banani, S. F., Russo, P. S., Jiang, Q. X., Nixon, B. T., & Rosen, M. K. (2012). Phase transitions in the assembly of multivalent signalling proteins. Nature, 483(7389), 336–340. https://doi.org/10.1038/nature10879

Liu, A. P., Aguet, F., Danuser, G., & Schmid, S. L. (2010). Local clustering of transferrin receptors promotes clathrin-coated pit initiation. J Cell Biol, 191(7), 1381–1393. https://doi.org/10.1083/jcb.201008117

Loerke, D., Mettlen, M., Yarar, D., Jaqaman, K., Jaqaman, H., Danuser, G., & Schmid, S. L. (2009). Cargo and dynamin regulate clathrin-coated pit maturation. PLoS Biol, 7(3), e57. https://doi.org/10.1371/journal.pbio.1000057

Malim, M. H., & Bieniasz, P. D. (2012). HIV Restriction Factors and Mechanisms of Evasion. Cold Spring Harb Perspect Med, 2(5), a006940. https://doi.org/10.1101/cshperspect.a006940

Manrique, S., Sauter, D., Horenkamp, F. A., Lulf, S., Yu, H., Hotter, D., Anand, K., Kirchhoff, F., & Geyer, M. (2017). Endocytic sorting motif interactions involved in Nef-mediated downmodulation of CD4 and CD3. Nat Commun, 8(1), 442. https://doi.org/10.1038/s41467-017-00481-z

Mariani, R., Kirchhoff, F., Greenough, T. C., Sullivan, J. L., Desrosiers, R. C., & Skowronski, J. (1996). High frequency of defective nef alleles in a long-term survivor with nonprogressive human immunodeficiency virus type 1 infection. J Virol, 70(11), 7752–7764. https://doi.org/10.1128/JVI.70.11.7752-7764.1996

Mariani, R., & Skowronski, J. (1993). CD4 down-regulation by nef alleles isolated from human immunodeficiency virus type 1-infected individuals. Proc Natl Acad Sci U S A, 90(12), 5549–5553. https://doi.org/10.1073/pnas.90.12.5549

Merrifield, C. J., Feldman, M. E., Wan, L., & Almers, W. (2002). Imaging actin and dynamin recruitment during invagination of single clathrin-coated pits. Nat Cell Biol, 4(9), 691–698. https://doi.org/10.1038/ncb837

Mettlen, M., Chen, P. H., Srinivasan, S., Danuser, G., & Schmid, S. L. (2018). Regulation of Clathrin-Mediated Endocytosis. Annu Rev Biochem, 87, 871–896. https://doi.org/10.1146/annurev-biochem-062917-012644

Pedersen, R. T. A., Hassinger, J. E., Marchando, P., & Drubin, D. G. (2020). Spatial regulation of clathrin-mediated endocytosis through position-dependent site maturation. J Cell Biol, 219(11). https://doi.org/10.1083/jcb.202002160

Pereira, E. A., & daSilva, L. L. (2016). HIV-1 Nef: Taking Control of Protein Trafficking. Traffic, 17(9), 976–996. https://doi.org/10.1111/tra.12412

Ren, X., Park, S. Y., Bonifacino, J. S., & Hurley, J. H. (2014). How HIV-1 Nef hijacks the AP-2 clathrin adaptor to downregulate CD4. Elife, 3, e01754. https://doi.org/10.7554/eLife.01754

Rosa, A., Chande, A., Ziglio, S., De Sanctis, V., Bertorelli, R., Goh, S. L., McCauley, S. M., Nowosielska, A., Antonarakis, S. E., Luban, J., Santoni, F. A., & Pizzato, M. (2015). HIV- 1 Nef promotes infection by excluding SERINC5 from virion incorporation. Nature, 526(7572), 212–217. https://doi.org/10.1038/nature15399

Saksela, K., Cheng, G., & Baltimore, D. (1995). Proline-rich (PxxP) motifs in HIV-1 Nef bind to SH3 domains of a subset of Src kinases and are required for the enhanced growth of Nef+ viruses but not for down-regulation of CD4. EMBO J, 14(3), 484–491. https://doi.org/10.1002/j.1460-2075.1995.tb07024.x

Schwenk, H. U., & Schneider, U. (1975). Cell cycle dependency of a T-cell marker on lymphoblasts. Blut, 31(5), 299–306. https://doi.org/10.1007/BF01634146

Serra-Moreno, R., Zimmermann, K., Stern, L. J., & Evans, D. T. (2013). Tetherin/BST-2 antagonism by Nef depends on a direct physical interaction between Nef and tetherin, and on clathrin-mediated endocytosis. PLoS Pathog, 9(7), e1003487. https://doi.org/10.1371/journal.ppat.1003487

Sun, Y., Leong, N. T., Jiang, T., Tangara, A., Darzacq, X., & Drubin, D. G. (2017). Switch-like Arp2/3 activation upon WASP and WIP recruitment to an apparent threshold level by multivalent linker proteins in vivo. Elife, 6. https://doi.org/10.7554/eLife.29140

Taylor, M. J., Perrais, D., & Merrifield, C. J. (2011). A high precision survey of the molecular dynamics of mammalian clathrin-mediated endocytosis. PLoS Biol, 9(3), e1000604. https://doi.org/10.1371/journal.pbio.1000604

Tokunaga, M., Imamoto, N., & Sakata-Sogawa, K. (2008). Highly inclined thin illumination enables clear single-molecule imaging in cells. Nat Methods, 5(2), 159–161. https://doi.org/10.1038/nmeth1171

Usami, Y., Wu, Y., & Gottlinger, H. G. (2015). SERINC3 and SERINC5 restrict HIV-1 infectivity and are counteracted by Nef. Nature, 526(7572), 218–223. https://doi.org/10.1038/nature15400

Wang, X., Chen, Z., Mettlen, M., Noh, J., Schmid, S. L., & Danuser, G. (2020). DASC, a sensitive classifier for measuring discrete early stages in clathrin-mediated endocytosis. Elife, 9. https://doi.org/10.7554/eLife.53686

Zhang, F., Wilson, S. J., Landford, W. C., Virgen, B., Gregory, D., Johnson, M. C., Munch, J., Kirchhoff, F., Bieniasz, P. D., & Hatziioannou, T. (2009). Nef proteins from simian immunodeficiency viruses are tetherin antagonists. Cell Host Microbe, 6(1), 54–67. https://doi.org/10.1016/j.chom.2009.05.008

