## Supplemental Figures for "Kinetic investigation reveals an HIV-1 Nef-dependent increase in AP-2 recruitment and productivity at endocytic sites"

**Figure S1.**

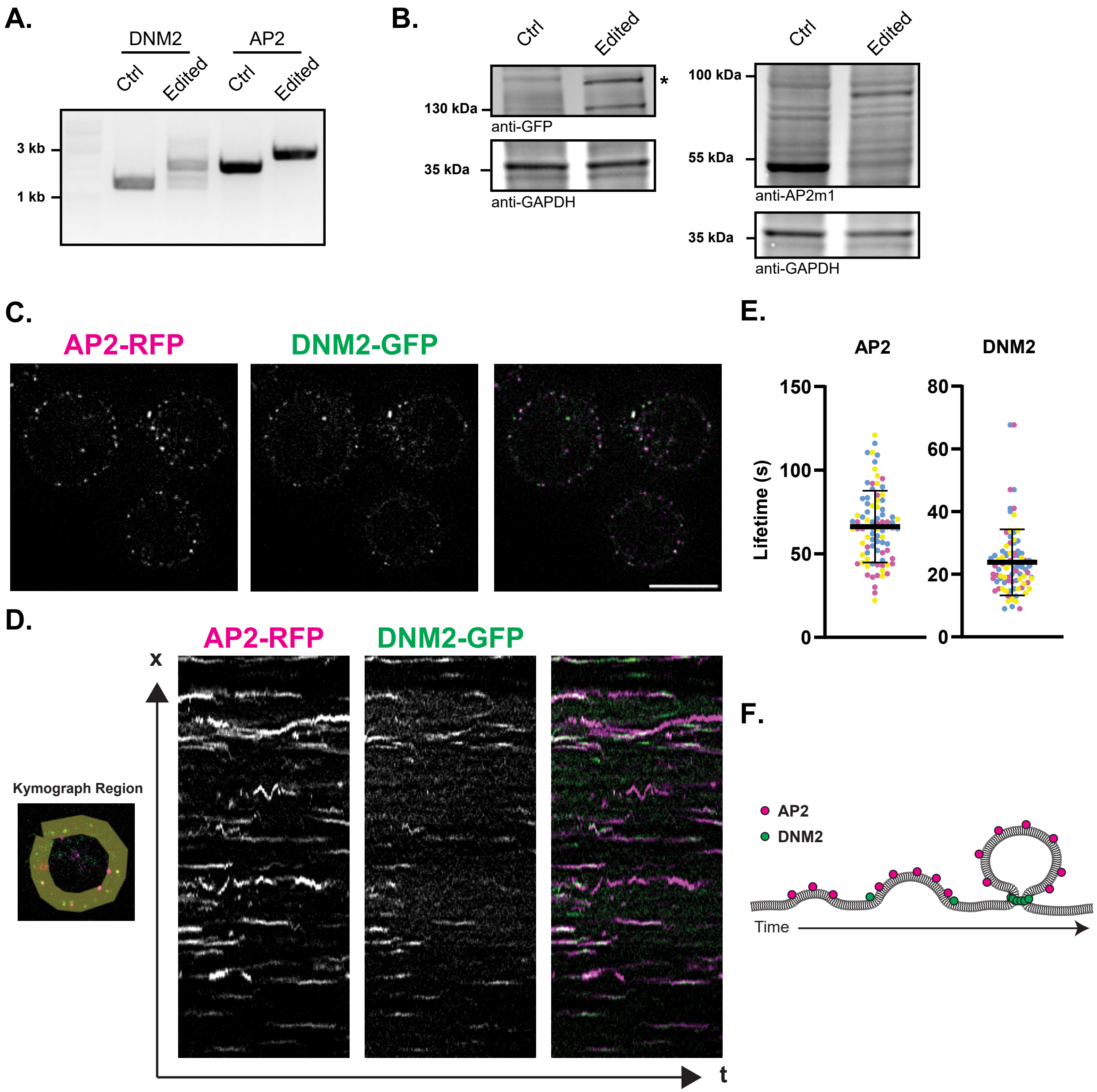
A.
DNM2
AP2
Ctrl
Edited
Ctrl
Edited
3 kb
1 kb
B.
Ctrl
Edited
*
130 kDa
anti-GFP
35 kDa
anti-GAPDH
Ctrl
Edited
100 kDa
55 kDa
anti-AP2m1
35 kDa
anti-GAPDH
C.
AP2-RFP
DNM2-GFP
D.
AP2-RFP
DNM2-GFP
x
t
Kymograph Region
E.
AP2
DNM2
Lifetime (s)
150
100
50
0
80
60
40
20
0
F.
AP2
DNM2
Time

**Supplementary Figure 1. AP-2-RFP DNM2-GFP dynamics in genome-edited Jurkat cells.** A. PCR analysis of genomic DNA from unedited (ctrl) and AP-2-RFP, DNM2-GFP edited (edited) Jurkat cells at the genomic loci of fluorophore insertion. B. Western blot of unedited (ctrl) and AP-2-RFP, DNM2-GFP edited (edited) Jurkat cells to confirm expression of fluorophore tagged protein at the expected molecular weight. Asterisk represents a specific, but unknown band. Samples were blotted with anti-GFP, anti-AP-2m1, and anti-GAPDH as the loading control. C. Midfocal plane image of Jurkat cells endogenously expressing AP-2-RFP and DNM2-GFP. Images were acquired through HiLo microscopy. Scale bar is 10 mm. D. Kymograph showing AP-2-RFP and DNM2-GFP dynamics over a 5 minute movie. E. Quantification of AP-2 and DNM2 lifetimes from kymographs. Each data point represents an endocytic site. The three colors represent different biological replicates. Since DNM2 sometimes shows multiple peaks within a single event, only the final peak was measured for lifetime determination.  F. Diagram illustrating internalization through the clathrin-mediated endocytosis (CME) pathway. The coat protein AP-2 is present for much of the internalization while most of the scission factor dynamin 2 (DNM2) is recruited as a rapid burst when the clathrin-coated vesicle is internalized.

**Figure S2.**

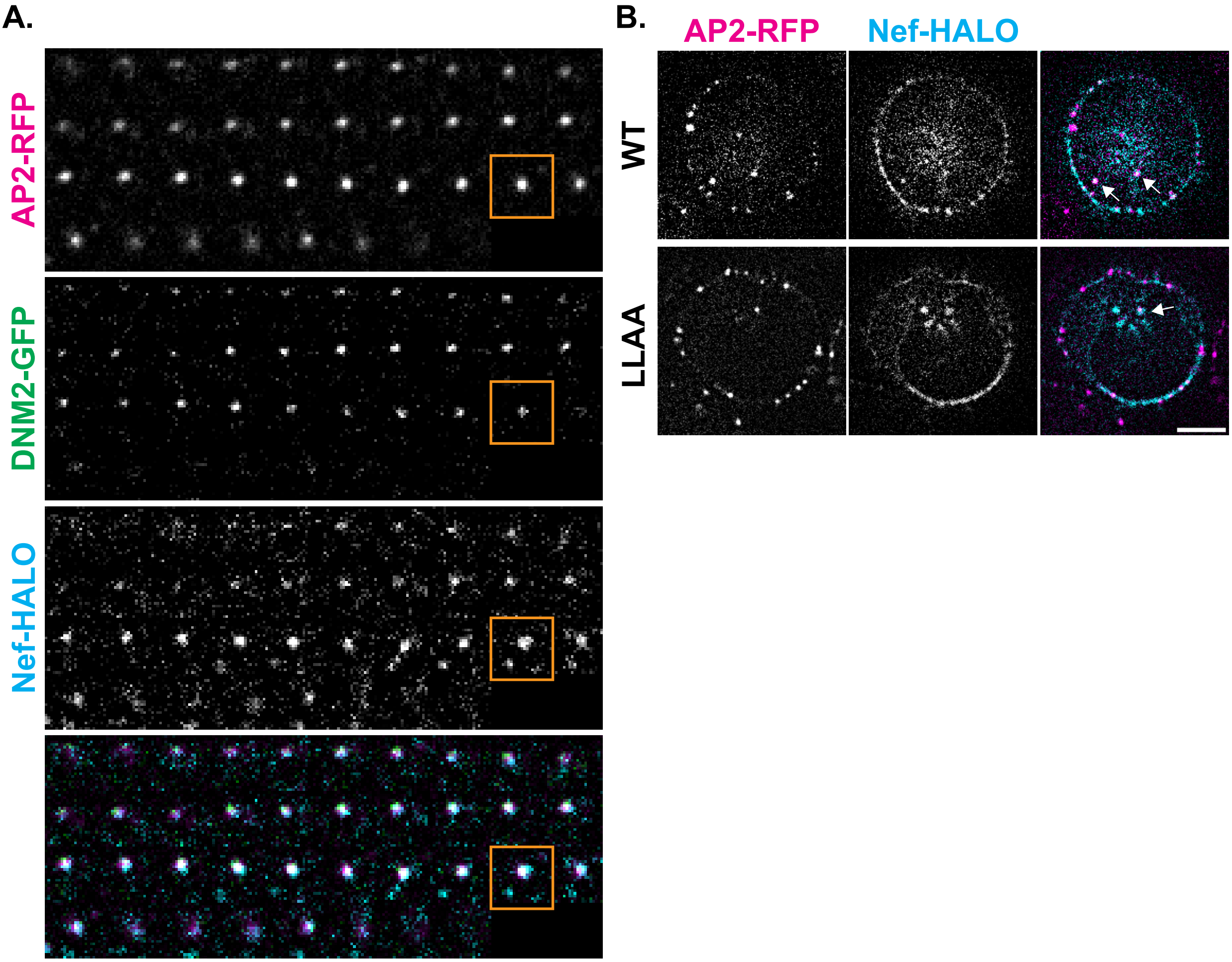

**Supplementary Figure 2. Nef is internalized during CME with the vesicle and colocalizes with intracellular AP-2 regardless of dileucine motif functionality.** A. Montage of a single Nef positive CME site in an AP-2-RFP and DNM2-GFP edited, Nef-HALO transfected Jurkat cell. Each frame in the montage is four seconds apart. The orange square indicates the frame before DNM2 signal rapidly decays. Shown in the bottom panel is a merge of the upper three channels. B. HiLo image midfocal plane of AP-2-RFP, DNM2 edited Jurkat cells transiently expressing WT or LLAA Nef-HALO. White arrows indicate intracellular AP-2 and Nef colocalization events. Scale bar is 5 µm.

**Figure S3.**

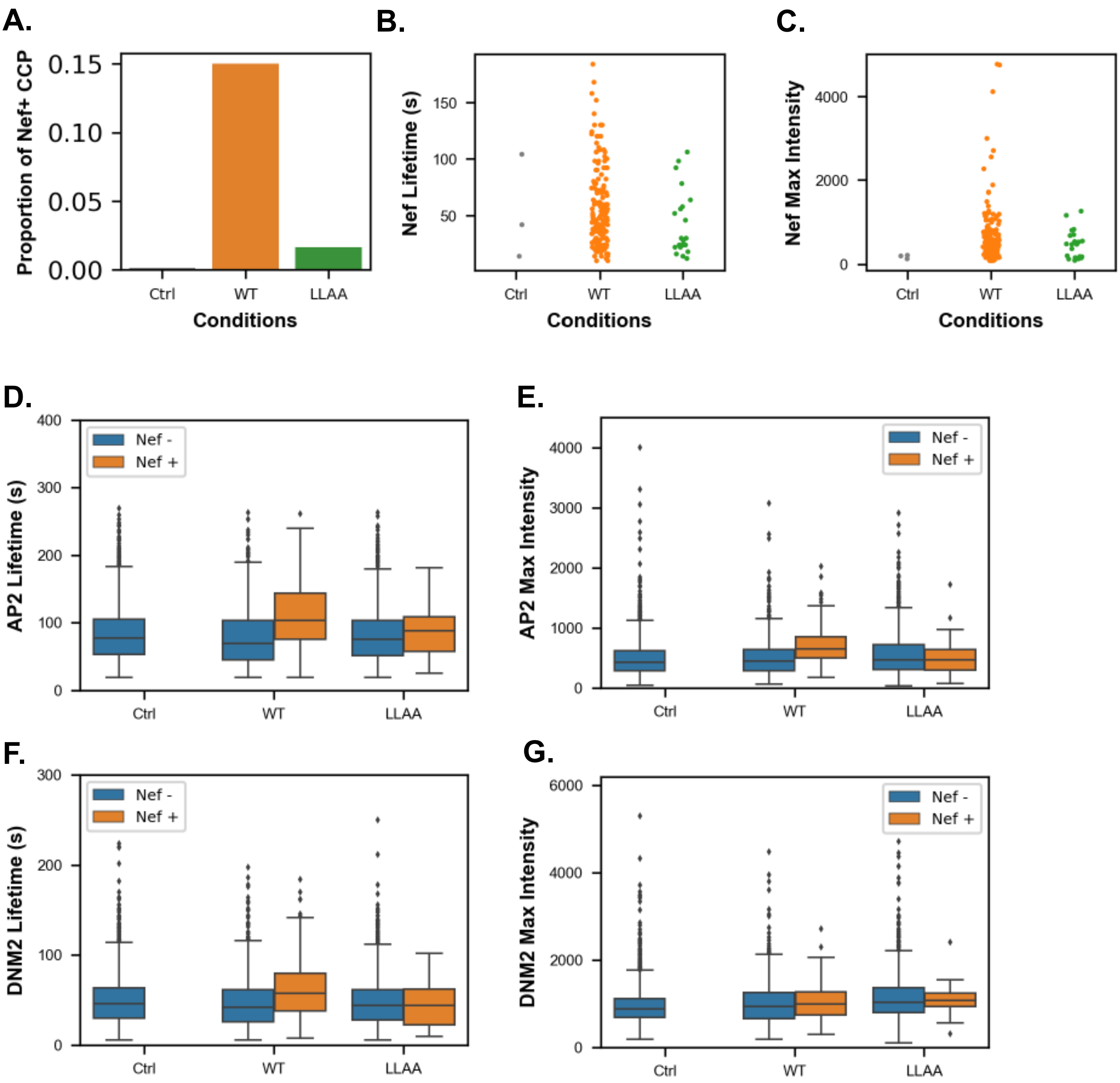
A.
B.
C.
Proportion of Nef+ CCP
0.15
0.10
0.05
0.00
Ctrl
WT
LLAA
Conditions
Nef Lifetime (s)
150
100
50
Nef Max Intensity
4000
2000
0
D.
E.
AP2 Lifetime (s)
400
300
200
100
Nef -
Nef +
AP2 Max Intensity
3000
1000
F.
G.
DNM2 Lifetime (s)
DNM2 Max Intensity
6000

**Supplementary Figure 3. Nef positive sites show dileucine dependent increase in AP-2 lifetime and recruitment.** A. Proportion of Nef+ CCP events out of all analyzed events in untransfected, WT Nef, and LLAA Nef transfected AP-2-RFP DNM2-GFP Jurkat cells. B. Nef lifetimes of detected Nef+ events in untransfected, WT Nef, and LLAA Nef transfected AP-2-RFP DNM2-GFP Jurkat cells. C. Nef max intensity of detected Nef+ events in untransfected, WT Nef, and LLAA Nef transfected AP-2-RFP DNM2-GFP Jurkat cells. D – F. Box and whisker plots of AP-2 lifetime (D), AP-2 Max intensity (E), DNM2 lifetime (F), and DNM2 Max Intensity (G). Events are from untransfected, WT Nef, and LLAA Nef transfected AP-2-RFP DNM2-GFP Jurkat cells. The blue box and orange box represent Nef- and Nef+ events respectively. Outliers are shown as black diamonds.

**Figure S4.**

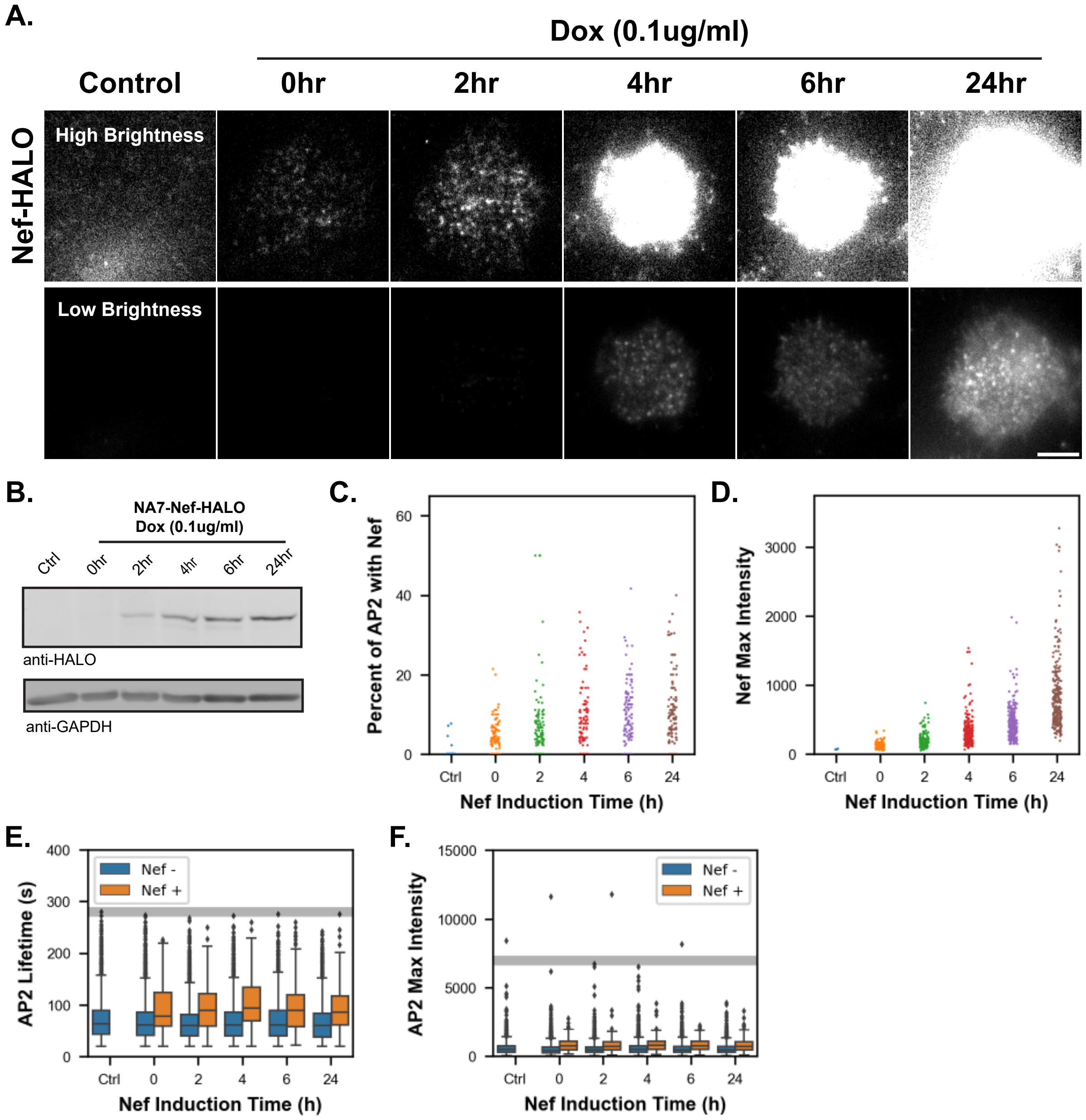
A.
Dox (0.1ug/ml)
Control
0hr
2hr
4hr
6hr
24hr
Nef-HALO
High Brightness
Low Brightness
B.
NA7-Nef-HALO
Dox (0.1ug/ml)
Ctrl
0hr
2hr
4hr
6hr
24hr
anti-HALO
anti-GAPDH
C.
Percent of AP2 with Nef
Nef Induction Time (h)
D.
Nef Max Intensity
Nef Induction Time (h)
E.
AP2 Lifetime (s)
Nef -
Nef +
Nef Induction Time (h)
F.
AP2 Max Intensity
Nef -
Nef +
Nef Induction Time (h)

**Supplementary Figure 4. NA7-Nef-HALO is inducibly expressed in AP-2-RFP, DNM2-GFP, NA7-Nef-HALO cell line.** A. TIR-FM images of Nef expression in either AP-2-RFP, DNM2-GFP Jurkat cells (control), or AP-2-RFP, DNM2-GFP, NA7-Nef-HALO Jurkat cells with indicated duration of doxycycline treatment. Top row of images were adjusted to show low range of Nef expression. Bottom row of images were adjusted to show high range of Nef expression. Scale bar is 5 μm. B. Western blot analysis of Nef expression in either AP-2-RFP, DNM2-GFP Jurkat cells (control), or AP-2-RFP, DNM2-GFP, NA7-Nef-HALO Jurkat cells with indicated duration of doxycycline treatment. NA7-Nef-HALO was detected with anti-HALO antibody. The loading control GAPDH was detected with anti-GAPDH antibody. C. Quantification of the percent of dynamic AP-2 sites that recruit Nef after different durations of Nef induction in the AP-2-RFP, DNM2-GFP, NA7-Nef-HALO Jurkat cell line. Each point represents a cell. D. Quantification of Nef max intensity at Nef positive spots after different durations of Nef induction. Each point represents a Nef+ CME site. E. Quantification of AP-2 lifetimes at Nef positive sites after different durations of Nef induction. Blue bars represent values from Nef-negative sites and orange bars represent values from Nef-positive sites. The blue bar for control sites refers to CME sites detected in the parental AP-2-RFP, DNM2-GFP Jurkat cell line. The gray bar indicates the estimated AP-2 lifetime corresponding to furthest outlier values in Nef- conditions. F. Quantification of AP-2 maximum intensity at Nef positive sites after different durations of Nef induction. Colored bars correspond to the same events as in E. The gray bar indicates the estimated AP-2 max intensity corresponding to furthest outlier values in Nef- conditions.
